# Modeling Adeno-Associated Viral Vector 6-mediated *In Vivo* Gene Delivery to Expanded Non-Mobilized Haemopoietic Stem Cells from Transfusion-dependent Thalassemia Patients in a Humanized Mouse

**DOI:** 10.1101/2025.06.17.659025

**Authors:** Janani Ramesh, Karthikeyan Kandasamy, Nur Nazneen Yusof, Ihsan Sujuandy, Choong Tat Keng, Liwei Chen, Bing Shao Chia, Min Liu, Zhisheng Her, Marek Kukumberg, Kian Chuan Sia, Siti Humairah Mohd Rodhi, Zhen Ying Fu, AJ Rufaihah, Alfredo Franco-Obregón, Mahesh Choolani, Binny Priya Sesurajan, Poh-San Lai, Shir Ying Lee, Pei Lin Koh, Qingfeng Chen, Shu-Uin Gan, Wei Leong Chew, Citra NZ Mattar

**Affiliations:** Experimental Fetal Medicine Group, Department of Obstetrics and Gynaecology, Yong Loo Lin School of Medicine, National University of Singapore, Singapore, Singapore; Genome Institute of Singapore, Agency for Science, Technology and Research, Singapore, Singapore; Institute of Molecular and Cell Biology, Agency for Science, Technology and Research, Singapore, Singapore; Healthy Longevity Translational Research Programme, Yong Loo Lin School of Medicine, National University of Singapore, Singapore; Phoenix Laboratory of Gene Therapy and Cell Therapy, Department of Surgery, Yong Loo Lin School of Medicine, National University of Singapore, Singapore, Singapore; Department of Surgery, Yong Loo Lin School of Medicine, National University of Singapore, Singapore, Singapore; School of Applied Sciences, Temasek Polytechnic, Singapore, Singapore; BICEPS Lab (Biolonic Currents Electromagnetic Pulsing Systems), National University of Singapore, Singapore, Singapore; Department of Obstetrics & Gynaecology, National University Hospital, National University Health Systems, Singapore, Singapore; Department of Obstetrics and Gynaecology, National University Centre for Women and Children (NUWoC), National University Health Systems, Singapore, Singapore; Department of Paediatrics, Yong Loo Lin School of Medicine, National University of Singapore, Singapore, Singapore; Department of Haematology-Oncology, National University Cancer Institute, National University Health Systems, Singapore, Singapore; Department of Laboratory Medicine, National University Hospital, Singapore, Singapore; Department of Paediatrics, Khoo Teck Puat – National University Children’s Medical Institute, National University Hospital, National University Health Systems, Singapore, Singapore; Synthetic Biology Translational Research Programme, Yong Loo Lin School of Medicine, National University of Singapore, Singapore, Singapore

**Keywords:** Hematopoietic stem cells, Humanized mouse model, Gene therapy, Adeno-associated virus, AAV, Dual-AAV6 transduction, β-thalassemia, HSC *in vitro* expansion

## Abstract

Hematopoietic stem cells (HSC) are important targets for gene modification therapies (GMT) as they originate several serious genetic conditions including the β-haemoglobinopathies. Potentially curative *ex vivo* GMT pose the barriers of accessibility, myeloablation-associated morbidity and prohibitive cost. *In vivo* GMT using non-integrating single-strand adeno-associated viral vectors (ssAAV) are a promising alternative that address these challenges directly, although the small ssAAV payload limits the capacity to package much larger gene or base editors. We investigated the feasibility of targeting human HSC *in vivo* with a dual-ssAAV6 strategy, which in future may be useful to deliver split-intein editing tools to overcome this limitation.

We engrafted NOD.Cg-Prkdcscid Il2rgtm1Wjl/SzJ humice with human hCD45^+^CD34^+^ HSC from transfusion-dependent β-thalassemic patients to test *in vivo* targeting of hCD45 cells with ssAAV6, then administered 5E+12 genomes/kg of ssAAV6-GFP/ssAAV6-mCherry. Humice showed peak single-transgene expression (GFP^+^ or mCh^+^) of 1.96-10.17%, and dual-transgene expression (GFP^+^mCh^+^) of 31.77% in circulating hCD45^+^ cells. Nested hCD45^+^ from liver, spleen and bone marrow showed single-and dual-transgene expression of 36.13-68.14% and 21.91-59.44% respectively. Secondary transplantation experiments demonstrated long-term persistence of AAV6-transduced hCD45 cells showing single-and dual-transgene expression of 9.19-60.72% and 7.15-9.19% respectively, with significant increase in expression from circulating cells. Minimal pro-inflammatory cytokine expression was observed following ssAAV6 administration in thalassemia humice compared with humice carrying non-thalassemia HSC.

Our model demonstrates the efficiency of *in vivo* ssAAV6-mediated targeting of thalassaemia HSC, potential long-term survivability of transduced cells, and feasibility of a dual-AAV strategy for gene editing, which offers a promising alternative to *ex vivo* GMT for β-haemoglobinopathies.

## INTRODUCTION

Hematopoietic stem cells (HSC) are fundamental to the maintenance of hematopoietic homeostasis, and genetic mutations affecting HSC function, including the haemoglobinopathies,^1^ primary immunodeficiencies (X-linked severe combined immunodeficiencies),^2^ neurometabolic and storage diseases (Gaucher disease, mucopolysaccharidoses),^3, 4^ congenital cytopenia (Fanconi anemia)^5^ and stem cell defects (e.g. Schwachman-Diamond syndrome)^6^ cause life-long relapsing morbidity. In these conditions, HSC produce structurally abnormal protein chains, severely restricted protein quantity, or are unable to differentiate into multiple lineages resulting in innate or adaptive immune cell deficiencies. Due to the multiplicity of critical metabolic, erythropoietic, regenerative, microenvironmental and protective functions provided by HSC, the collective medical and socioeconomic burdens of this group of genetic diseases are substantial.^7^

HSC-directed gene modification therapies (GMTs) carry as-yet-unrealized therapeutic potential for these monogenic diseases, and are challenging due to numerous physiological barriers.^8, 9^ *Ex vivo* autologous HSC-directed GMT is currently the preferred approach, as precise genetic correction without undesirable off-target mutations can be assessed, and gene-modified CD34^+^ HSC can be expanded, if necessary, prior to transplantation. Zynteglo®, Lyfgenia® and Casgevy® have all been approved by the US Food and Drug Administration for treatment of transfusion-dependent β-thalassemia and sickle cell disease.^10^ *Ex vivo* GMT can achieve high levels of *in vitro* editing or transduction, but still depends on successful engraftment and proliferation of modified cells; current data suggest that modified HSC are capable of this and contribute substantially to reversal of phenotype.^11^ While these therapeutic breakthroughs are major steps towards providing cures for all, the downsides are substantial, including conditioning-related morbidity, graft failure, graft-v-host disease (GVHD) and opportunistic infections.^12^ Because of highly-specialized current good manufacturing practice (cGMP) infrastructure requirements and extensive conditioning of recipients prior to administrating these cell products, accessibility and affordability are barriers.^13^ *In vivo* HSC-directed GMTs may eventually circumvent these barriers and reach a wider spectrum of recipients, but also raise concerns about safety and efficiency. Integrating lentiviral vectors, as currently used for *ex vivo* modification, cannot be used *in vivo* due to mutagenesis, while “safer” non-pathogenic adeno-associated viral vectors (AAV)^14^ that are vehicles for the majority of systemically-administered *in vivo* GMTs (e.g. Hemgenix®, Roctavian® for haemophilia) have payloads too small (<4.7 kb) to accommodate large transgenes or gene editors (9– 19 kb).^15^ To overcome this, GMT tools can be split and packaged into two separate AAV vectors, requiring dual transduction of target cells by both AAVs.^16^

HSC are maintained in a rich reservoir and are tightly regulated within the hematopoietic niches to balance self-renewal and differentiation, to sustain haemopoiesis.^17^ Additionally, adult HSC are rare in peripheral blood and require efficient vectors to achieve a sufficient transduction in these quiescent cells.^8^ Mobilization of niche HSC with granulocyte colony stimulating growth factor (GCSF) transiently enriches circulating peripheral blood HSC (PBHSC) and may enhance *in vivo* HSC transduction,^18, 19^ is relatively safe and widely used, but may rarely cause bone pain, pulmonary embolism and splenic rupture.^20^ Thus, targeting these elusive cells is a particular challenge of *in vivo* GMT.

In preparation for proof-of-concept *in vivo* AAV6-mediated genetic editing of common β-globin mutations, we performed a feasibility study of dual-AAV6 targeting of human CD34^+^ HSC in a humanized mouse model. Modelling *in vivo* HSC-directed GMT with preclinical models is challenging. Knock-in/knock-out mouse models mimicking HSC-related genetic disorders are valuable for proof-of-concept data, while NHP data is informative for long-term durability and safety.^21, 22^ Despite overlapping features, inherent species-specific physiological differences may render clinical vectors less effective in these models.^23^ Adult human HSC carrying pathological mutations show functional profiles distinct from non-diseased HSC.^24^ While there is no ideal model, efficiency of and barriers to *in vivo* GMT can be revealed in humanized mice (humice) produced with human adult HSC. We have produced such a model engrafted with transfusion-dependent β-thalassemia (TDT) patient-derived HSC to interrogate the efficiency of dual AAV6-mediated HSC transduction. Our model was transplanted with *in vitro* expanded non-mobilized PBHSC as none of our patients underwent medically indicated mobilization at the time of sample collection. We used single-stranded ssAAV6 capable of carrying a larger payload than self-complementary AAV, as a clinical strategy applicable to *in vivo* GMT (**Figure 1**). Our study demonstrates the feasibility of generating patient-specific humice with non-mobilized thalassaemic HSC, circumventing the need for GCSF mobilization, which is often contraindicated or risky in patients with haemoglobinopathies. The model achieves substantial expansion and engraftment of human HSC, enabling in vivo assessment of gene targeting outcomes, and directly addresses the clinical need for accessible and scalable preclinical platforms to predict patient-specific responses.

**Figure 1.**
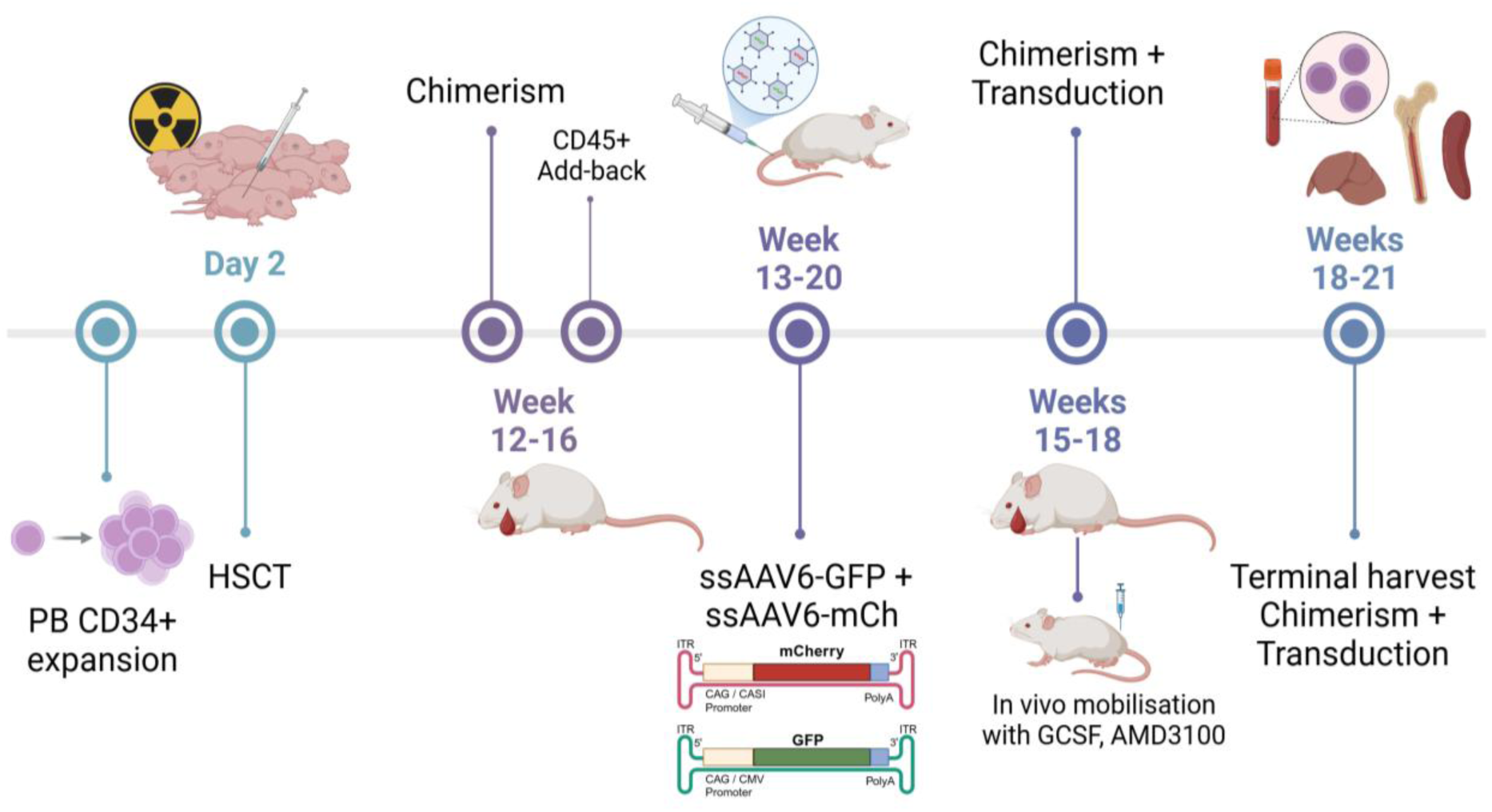
Experimental design for expansion and injection of CD34+ HSC into irradiated NOD-SCID-IL2rgnull (NSG) mice to generate humanized mice (humice), showing add-back of peripheral blood mononuclear cells in humice with low engraftment, administration of AAV6-CASI-GFP and AAV6-CMV-mCh or AAV6-CAG-GFP and AAV6-CAG-mCh, and analyses of outcomes following in vivo transduction.

## METHODS

### Hematopoietic isolation, expansion and characterization

Pediatric and adult patients previously diagnosed with TDT attending the National University Hospital, Singapore, were consented under DSRB 2021/00783, and healthy volunteers were consented under NUS-IRB-2021-424. Briefly, non-mobilized peripheral blood (PB) collected via venipuncture into EDTA tubes was diluted with equal volume of EasySep buffer (StemCell Technologies, Vancouver, Canada) containing 2% antibiotic-antimycotic (Anti-Anti) (Gibco, Grand Island, NY) and overlayed on Ficoll-Paque Plus (Cytiva™, Marlborough, MA, USA), centrifuged (brakeless) at 400G for 20min, and the buffy coat collected. CD34^+^ cells (nmPBHSC) were isolated from mononuclear cells using EasySep™ Human CD34 Positive Selection Kit II (StemCell Technologies) following manufacturer’s instructions, with 150μL/mL RapidSpheres™ and 200μL/mL antibody. Isolated nmPBHSC were washed and plated at 1E+5cells/mL in StemSpan SFEM II media (StemCell Technologies) containing SCF, TPO, FLT3L, SR1, VEGF, vitamin C, IL-3, IL-6 for 14 days, then characterized by flow cytometry. Mobilized PBHSC (LONZA Group AG, Basel, Switzerland) were cultured similarly.

Single-cell suspensions of expanded PBHSC were analyzed for HSC surface markers using appropriate antibodies (**Supplemental Table S1**) as previously reported.^25^ 5E+3 PBHSC were resuspended in 300μL of Iscove’s DMEM with 5% FBS (Gibco), suspended in 3mL MethoCult^TM^ SF H4636 (StemCell Technologies) following manufacturer’s instructions, cultured at 37°C, 5% CO2 for 14d prior to colony counting. Images were taken with an EVOS M7000 Imaging System (Thermo Fisher Scientific Inc., Waltham, MA) at 4X and 10X magnification.

### Single-strand AAV6 production with various promoters and *in vitro* transduction of HSC

AAV6-CASI-GFP and AAV6-CMV-mCherry were constructed at the Vector Core@Genome Institute of Singapore (A*STAR) with pAAV-Rep1Cap6 plasmid (from cloning in AAV6 capsid gene (GenBank AF028704.1) fragment into the pAAV-RepCap plasmid backbone using Gibson Assembly) as previously described.^16^ Briefly, AAVs were packaged via triple transfection of 293AAV cell line (AAV-100, Cell Biolabs Inc, San Diego, CA) plated in HYPERFlask ‘M’ (Corning Inc, Somerville, MA) in growth media consisting of DMEM, glutaMax, pyruvate with 10% foetal bovine serum (FBS) (Thermo Fisher Scientific Inc.), supplemented with 1XMEM non-essential amino acids (Gibco). Transfection was performed at 70–90% confluence with 200 μg pHelper (Cell Biolabs Inc), 100 μg pAAV-Rep1Cap6, 100 μg pZac-CASI-GFP or 100 μg pAAV-CMV-mCherry-NLS (#165442, Addgene, Watertown, MA) mixed in 5ml DMEM and 2 mg PEI “MAX” (Polysciences, Warrington, PA) at a PEI: DNA mass ratio of 5:1. Media was amended to 2% FBS one day after transfection, cells harvested 48–72h later, freeze-thawed 3x, supernatant collected after centrifugation at 4000Gx5min, and treated with 50U/ml Benzonase (Sigma-Aldrich, St Louis, MO), 1 U/ml RNase cocktail (Invitrogen, Carlsbad, CA) for 30min at 37°C to remove unpackaged nucleic acids. Lysate was loaded on a discontinuous density gradient (6mL each of 15%, 25%, 40%, 60% Optiprep (Sigma-Aldrich) in 29.9mL Optiseal polypropylene tube (Beckman-Coulter, Pasadena, CA), ultracentrifuged at 54000rpm, the 40% fraction extracted and dialyzed using Amicon Ultra-15 (100kDa MWCO, Merck Millipore, Darmstadt, Germany), and purified AAV titered by real-time qPCR with ITR-sequence-specific primers (**Supplemental Table S2**).^26^

AAV6-CAG-GFP and AAV6-CAG-mCherry were produced at the Phoenix Vector Core Facility NUS, by 72h double-transfection of AAV293 (Gibco) with pCAG-GFP expression plasmid (Addgene Plasmid #139980) or an in-house cloned pAAV-CAG-mCherry plasmid, and helper and capsid plasmid pDGM6 (Addgene Plasmid #110660). AAV6-CAG-GFP and AAV6-CAG-mCherry were generated using AAV-MAX Helper-Free AAV Production System Kit (Gibco), as previously described.^27^ Briefly, cell lysates were treated with 90 U/mL Universal Nuclease (Pierce Biotechnology Inc., Waltham, MA), purified via iodixanol gradient ultracentrifugation (Optiprep™, Progen, Heidelberg, Germany) at 55000rpm, AAV particles concentrated with Amicon Ultra-4 (100 kDa MWCO; Merck Millipore), and qPCR-titered using primers specific for CAG (**Supplemental Table S2**).

*In vitro* transduction of HSC was assessed for all ssAAV6 vectors at 1E+5 and 1E+6 multiplicities of infection (MOI), on mPBHSC (LONZA) on 24-well plates (1E+5 cells/well) in PBHSC media at 37°C, 5%CO2 for 24h. Cells were washed, replated in new media, incubated for another 72h (96h total incubation), and stained with Sytox Blue (L/D) for flow cytometry.

### Humanized mouse model, secondary transplantation and ssAAV6 injection

Time-mated female NOD-SCID-IL2rg*null* (NSG) mice purchased from Jackson Laboratory (Bar Harbor, Maine) were housed in pathogen-free conditions at the Biological Resource Centre, Agency for Science, Technology and Research, Singapore (A*STAR) with controlled 12-hour light-dark cycle, with ad libitum access to food. All experiments and procedures were approved by A*STAR Institutional Animal Care and Use Committee (IACUC) in accordance with the guidelines of the Agri-Food and Veterinary Authority and the National Advisory Committee for Laboratory Animal Research (NACLAR) of Singapore. 1-3d old neonates were sublethally irradiated with 1Gy and injected intrahepatically with 4.5E+5 human CD34^+^ nmPBHSC (thalassemia or non-thalassemia) or mPBHSC, creating three experimental groups: Thal-HM (engrafted with non-mobilized PBHSC from TDT patients), Normal-HM (engrafted with non-mobilized PBHSC from non-thalassemia volunteers) and Mob-HM (engrafted with GCSF-mobilized PBHSC). Humice were evaluated at 12w post-transplantation for immune reconstitution by human hCD45^+^ cells by flow cytometry and reassessed serially if the initial chimerism was low.

hCD45^+^ cells were isolated from PB, bone marrow (BM), liver and spleen of Thal-HM (from two TDT donors) sacrificed at 14w using EasySep™ Release Human CD45 Positive Selection Kit (StemCell Technologies) following manufacturer’s instructions and used for secondary transplantation, with 1.5E+6 cells injected intrahepatically into irradiated neonatal NSG mice. PB, BM, liver and spleen were analyzed at 14w for long-term repopulating hCD45. Humice with hCD45^+^ >2% were injected intravenously with 5E+12 vector genome (vg)/kg of ssAAV6-GFP, ssAAV6-mCh or a 1:1 mixture of both, at 12w-20w post-transplantation. Humice with initial chimerism <2% received intravenous add-back of 5E+6 CD34^-^ mononuclear cells from the same donor source after CD34^+^ harvest to boost hCD45 chimerism, reassessed and injected with ssAAV6 2w later. Transgene expression was serially monitored from 96h-4w, when PB serum and cells, BM, liver and spleen were harvested for analyses. Outcomes compared among the three experimental groups were immunotoxicity, viz development of graft-v-host disease (GVHD), or ssAAV6 injection (AAV^+^ or AAV^-^). GVHD was induced in Normal-HM by injection of pooled nmPBHSC from various donors, and was clinically diagnosed (alopecia, hunching, lethargy, weight loss).

### Analyses of AAV transduction and cytokine response

Single-cell suspensions of expanded PBHSC and recovered human and murine CD45^+^ cells were incubated with FC block, appropriate antibodies (**Supplemental Table S1**), assayed using X-20 BD Fortessa flow cytometer, raw data compensated with single-color controls and readouts analyzed using FlowJo (BD Biosciences, East Rutherford, NJ, USA) as previously reported.^25^

Vector copy number (VCN) was assessed by qPCR as previously described.^28^ Briefly, genomic DNA (15ng) extracted with Qiagen DNeasy Blood and Tissue Kits for DNA Isolation (Maryland, USA) was subjected to a 20µL PCR reaction (FastStart SYBR Green Master, Roche, Germany) using forward (5′-GGAACCCCTAGTGATGGAGTT-3′) and reverse (5′-CGGCCTCAGTGAGCGA-3′) primers amplifying the ITR sequence (annealing temperature, 60°C). Equivalent loading was verified using forward (5′-ACCCGTTGACTCCGACTT-3′) and reverse (5′-ACCACTAGGCGCTCACTGTTC-3′) primers to amplify a 112bp region of the human GAPDH gene. VCN was expressed per diploid genome (6.6pg DNA), and the calculated limit of detection was 1 vector genome (vg) per 227 diploid genomes (1vg in 34 cells). To estimate VCN in human cells, the ratio of GFP expression in human hCD45 to GFP expression in mouse and human CD45 was calculated and multiplied to total VCN. Naïve DNA from non-transduced mice served as negative controls.

Human-and mouse-specific cellular responses were analyzed from terminal harvest plasma using high-throughput antibody-based multiplex immunoassays with magnetic beads, employing preconfigured panels specific for human (Procartaplex, Human Th1/Th2 Cytokine Panel 11plex) and mouse species (Procartaplex, Mouse Th1/Th2 Cytokine Panel 11plex, both ThermoFisher Scientific) on the Luminex platform (Luminex Corporation, Austin, TX, USA). Antibodies used in the species-specific kits were raised against an epitope of the analyte protein of that species. Briefly, samples and standards for 11 target analytes were incubated in turn with antibody magnetic beads (18h, 4°C), detection antibodies (30min) and Streptavidin-Phycoerythrin (30min), washing between steps. All incubations were performed on an orbital shaker (600rpm). Median immunofluorescence was measured on the Luminex 200 using xPONENT® 4.0 software (Luminex Corporation, Austin, TX, USA) to analyze selected cytokines (**Supplemental Table S3**). Analyte concentrations were adjusted based on a dilution factor of 2. Standard curves were generated with a 5-parameter logistic algorithm, reporting values for both mean fluorescence intensity (MFI) and extrapolated analyte concentration data. Changes in cytokine expression were compared by Z-score (±2SD) or by fold-change from the median cytokine level in control humice (**Figure 5A-D**), which were Normal-HM without GVHD. For Z-score values, cytokine data was preprocessed by converting values below the lower limit of detection (LLOD) reported as “<X” into numeric values dividing the detection limit by 2, reducing the impact of undetectable values. The baseline group of control humice (Normal-HM without GVHD) was used as the reference, with mean cytokine values calculated separately for humans and mice. Fold changes were determined relative to the baseline, and Z-score normalization was applied across each cytokine row to standardize the data for comparative purposes. Human and mice data are processed independently to maintain species-specific insights.

Anti-AAV6 IgG ELISA (Creative Diagnostics, Shirley, NY, USA) was performed following the manufacturer’s instructions. Briefly, serum extracted from humice at sacrifice was diluted 1:50 and added to microwells precoated with AAV6 capsid protein. After incubation and washing, enzyme-linked polyclonal anti-human IgG antibody was added. ELISA development was performed using TMB substrate, and the reaction was terminated with stop reagents. Absorbance was measured at 450/620 nm using a Biotek Epoch Microplate Spectrophotometer (Agilent Technologies, Santa Clara, CA, USA). Mean absorbance (OD) was calculated, and results were expressed in arbitrary units (U) using the formula: Units (U) = (Avg OD) / (Cut-off), where the cut-off value was determined from the manufacturer-provided cut-off control.

### Statistical analysis

Ordinary one-way and 2way ANOVA followed by Tukey or Šídák’s multiple comparisons test, and multiple unpaired parametric t tests correcting for multiple comparisons using Holm-Šídák method, were performed using GraphPad Prism version 10.4.0 for Windows, GraphPad Software, Boston, Massachusetts USA, www.graphpad.com. Data are represented as superimposed bars where relevant. p<0.05 was taken as significant.

## RESULTS

### Expansion of non-mobilized peripheral blood hematopoietic stem cells to assess AAV6 transduction efficiency

Non-mobilized peripheral blood HSC (PBHSC) from patients with TDT (Thal-PBHSC) and healthy persons without thalassemia (Normal-PBHSC) demonstrated cell number fold-increases of 3.93±2.99 and 3.43±1.42 respectively following *in vitro* expansion with stem cell growth factors for 14 days (d14), peaking between d8 and d14 (differences not significant (ns), **Figure 2A**). HSC were characterised via expression of hematopoietic stem and progenitor cell (HSPC) markers - long-term HSC (Lin-CD34+CD90+CD49f+), short-term HSC (Lin-CD34+CD90-CD38-CD49f+) and hematopoietic progenitor cells (Lin-CD34+CD90-CD38+CD49f+), assessed by flow cytometry (**Figure 2B**). There were higher proportions of HSPC in expanded Normal-PBHSC than in Thal-PBHSC by d8 (80.22±9.80%, (n=9), vs. 55.16±16.30% (n=5), p<0.005) and d11 (83.52±6.09% vs 42.69±15.14%, p<0.0001) while total CD34^+^ (57.95±8.26% vs. 56.08±18.15%), LT-HSC (12.50±6.45% vs. 5.39±5.80%) and ST-HSC (9.72±3.58% vs. 2.49±1.56%) were similar by d14 (**Figure 2C**). Though absolute cells counts expanded in the range of 20.94E+3–8332.64E+3 cells/mL by d14 in both samples, increases in mononuclear cells (8332.64±5845.90 E+3 vs. 449.99±489.35 E+3/mL, p<0.005) and ST-HSC (440.64±160.22E+3 vs. 24.85±16.66E+3/mL, p<0.001) were greater in Thal-PBHSC (**Figure 2D**). Both sources showed propensity to differentiate along erythroid lineages following expansion; we observed similar numbers of post-culture erythroid Burst-forming (BFU-E) and Colony-forming (CFU-E) erythroid progenitors, granulocyte-macrophage (CFU-GM) and common myeloid (granulocyte, erythrocyte, monocyte, megakaryocyte, CFU-GEMM) progenitors in both groups (**Figure 2E**); CFU-GM was the most frequently observed in Thal (99.83 units) and Normal (85.17 units) samples. For transplantation of NOD-SCID-IL2rg^null^ (NSG) mice, Thal-PBHSC donor inoculum (n=3) comprised 98.62±1.03% Lin-negative, 33.04±7.37% CD34^+^ cells, 1.26% long-term LT-HSC, 17.41% short-term ST-HSC and 14.36% HSPC, while Normal-PBHSC (n=5) comprised 82.65±23.30% Lin-negative, 22.50±27.56% CD34^+^ cells, 3.69% LT-HSC, 0.48% ST-HSC and 8.92% HSPC. Mobilized human PBHSC from non-thalassemia donors treated with granulocyte colony-stimulating factor (mPBHSC) prior to collection comprised 76.78±32.45% CD34^+^, 0.25±0.28% LT-HSC, 25.43±30.06% ST-HSC and 52.54±39.99% HSPC, and were not expanded *in vitro*.

**Figure 2.**
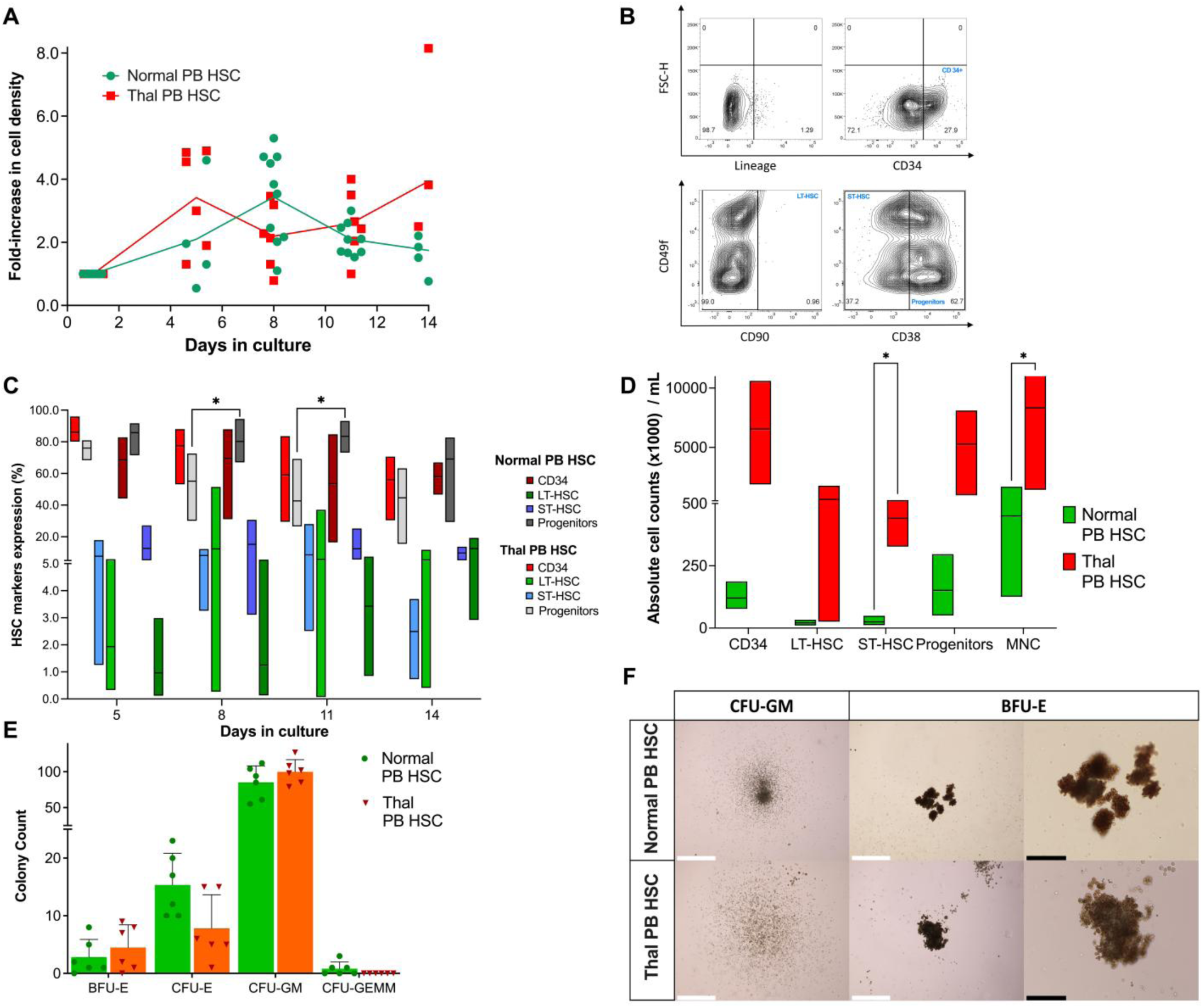
In vitro expansion of hematopoietic stem cells (HSC). **A** Fold-increase in cell density between non-thalassemia (Normal) and thalassemia (Thal) PBHSC over a 14-day culture. **B** Representative flow cytometry gating characterizing CD34+ HSC into long-term HSC (Lin^-^CD34^+^CD90^+^CD49f^+^), short-term HSC (Lin^-^CD34^+^CD90^-^CD38^-^CD49f^+^) and hematopoietic progenitor cells (Lin^-^CD34^+^CD90^-^CD38^+^CD49f^+^). **C** HSC surface marker expression in culture-expanded Normal and Thal PBHSC after 14 days, by flow cytometry. **D** Absolute cell counts of HSC subsets in Normal and Thal PBHSC after culture-expansion. **E** Absolute counts of Normal and Thal PBHSC-derived colonies in MethoCult SF H4636 media and their representative images. **F** White scale bar represents 650µm (4x magnification) and black scale bar represents 275µm (10x magnification). *p<0.05, **p<0.01, ***p<0.001, ****p<0.0001. PBHSC – peripheral blood hematopoietic stem cells.

### Chimerism in humanized mice of expanded non-mobilized PBHSC and secondary transplantation

Knowing the challenges of CD34^+^ HSC engraftment and human hematopoiesis recapitulation in humice, largely due to post-transplantation HSC differentiation,^29^ we quantitatively assessed circulating (in peripheral blood) and nested (in bone marrow (BM), liver and spleen) human CD45^+^ cells (hCD45) and CD34^+^ subsets in humice from 12w post-transplantation (**Figure 3A**). Circulating hCD45 chimerism in Thal-HM ranged from 4.15±7.21% at 12w post-transplantation (n=17) to a peak of 10.34±10.66% at 14w (n=12) before becoming almost undetectable at 20w (0.10±0.06%, n=8). Normal-HM required earlier assessment and terminal harvest due to evolving graft-v-host disease (GVHD), evidenced by ruffling and loss of fur, hunching and lethargy from 6w. hCD45 chimerism in peripheral blood was 60.34±33.20% at 6w and 38.06±30.73% at 9w, eventually decreasing to 11.445±15.73% by 10w at sacrifice (n=17), significantly higher at 6w and 9w than in Thal-HM and mPB-HM at all timepoints. Peripheral blood engraftment of mPBHSC resulted in hCD45 chimerism of 6.806±18.28% at 12w (n=16), peaking at 9.08±7.99% by 20w (n=6, **Figure 3B**); nested hCD45 chimerism ranged from 4.55-44.06% (data not shown). Where detectable in recovered circulating hCD45, CD34^+^ cell counts were low, ∼0.02-0.03% in Normal-HM and mPB-HM (n=8).

**Figure 3.**
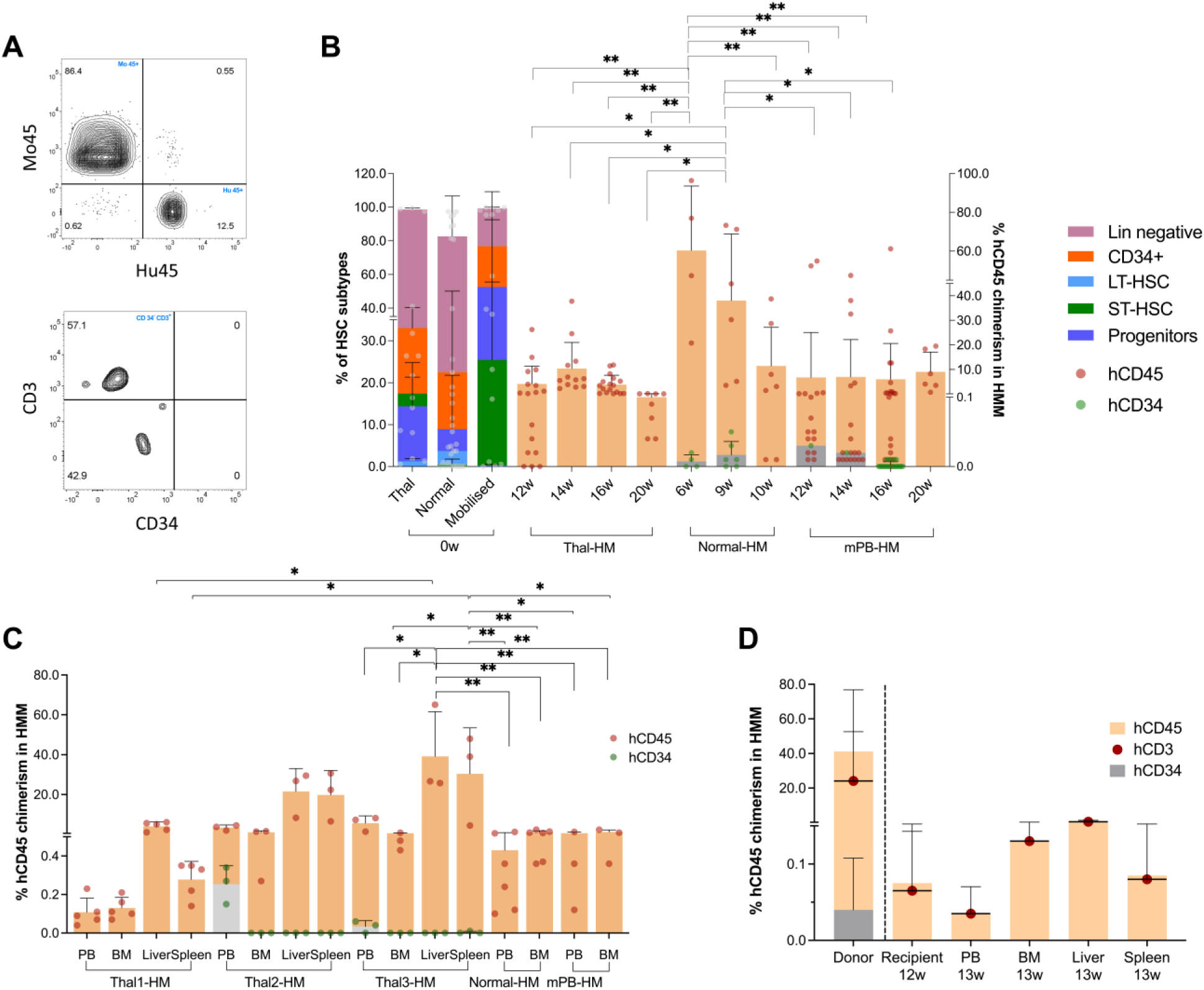
Primary and secondary transplantation of expanded hematopoietic stem cells in NSG mice. **A** Representative flow cytometry gating to assess human chimerism following transplantation and to determine frequencies of hCD34 and hCD3 pre-gated from hCD45+ in a humice model. **B** Pre-transplantation cell subset composition of Thal and Normal PBHSC and non-expanded mobilized mPBHSC and human hCD45 chimerism in Thal-HM, Normal-HM and mPB-HM measured over weeks. **C** Chimerism of hCD45 nested in organs at terminal harvest in Thal-HM, Normal-HM and mPB-HM. **D** Secondary transplantation of human CD45+ cells harvested from primary Thal-HM at 14w, showing composition of hCD45, hCD3 and hCD34 in retrieved “donor” cells, and secondary engraftment at 12w in PB and organs. *p<0.05, **p<0.01, ***p<0.001, ****p<0.0001. Thal-HM – humanized mice engrafted with expanded thalassemia HSC; Normal-HM – humanized mice engrafted with expanded non-thalassemia HSC; mPB-HM – humanized mice engrafted with mobilized, non-expanded non-thalassemia HSC.

Thal-HM showed circulating and nested hCD45 chimerism levels ranging from 0.1-65.07% by 21w at harvest, with the highest chimerism observed in Thal3-HM (39.17±22.43% in liver, 30.47±22.30% in spleen, exceeding levels in PB at 5.61±3.71% and BM at 0.57± 0.21%, p<0.01), significantly higher than chimerism in Thal1-HM (liver 4.01±2.22% and spleen 0.28±0.09%, p<0.01). BM chimerism was lowest among harvested organs across all Thal-HM groups at 0.1-1.55%. Normal-HM showed PB and BM chimerism of 0.43±0.42% and 1.01±0.76% respectively (harvested at 10w, due to GVHD), similar to mPB-HM which showed 0.57–1.20% chimerism in PB and BM at the 21w sacrifice (n=3, **Figure 3C**).

hCD45 were significantly higher in Thal3-HM liver and spleen compared to PB and BM of Thal3-HM, Normal-HM and mPB-HM. CD34^+^ cells were detected in Thal2-HM and Thal3-HM in PB (n=6), accounting for 0.03-0.25% of hCD45.

To boost low hCD45 chimerism in primary humice in preparation for *in vivo* AAV6 transduction and secondary transplantation, we added back 5E+6 hCD45 to two Thal-HM (Thal2-HM and Thal3-HM, both carrying IVS2-654C>T mutation) from the original donor PBHSC. Post-addback hCD45 reached 28.20-38.94% after 2-3w, and 41.20±35.61% at terminal harvest. Isolated primary hCD45 comprising 69.57±39.88% CD3^+^ and 0.34±0.63% CD34^+^ cells from these humice were transplanted into secondary NSG recipients. 13w later, secondary hCD45 chimerism was detected at 0.04±0.03% in PB (91.82±8.63% CD3^+^), and 0.58±0.78% in liver (81.64±14.43% CD3^+^); no hCD34^+^ cells were detected (**Figure 3D**).

### *In vitro* and *in vivo* single and dual AAV6 transduction of human HSC

We performed *in vitro* and *in vivo* transduction with two AAV6 vector pairs, AAV6-CAG-GFP/AAV6-CAG-mCh, and AAV6-CASI-GFP/AAV6-CMV-mCh, and transgene expression was quantified via flow cytometry (**Figure 4A**). *In vitro* transduction produced dual GFP^+^mCh^+^ expression reaching 25.39±1.54% at 1000E+3 MOI of AAV6-CAG-GFP and AAV6-CAG-mCh (**Figure 4B**); single and dual transduction were both higher with CAG promoters at 100E+3 and 1000E+3 MOI (p<0.003).

**Figure 4.**
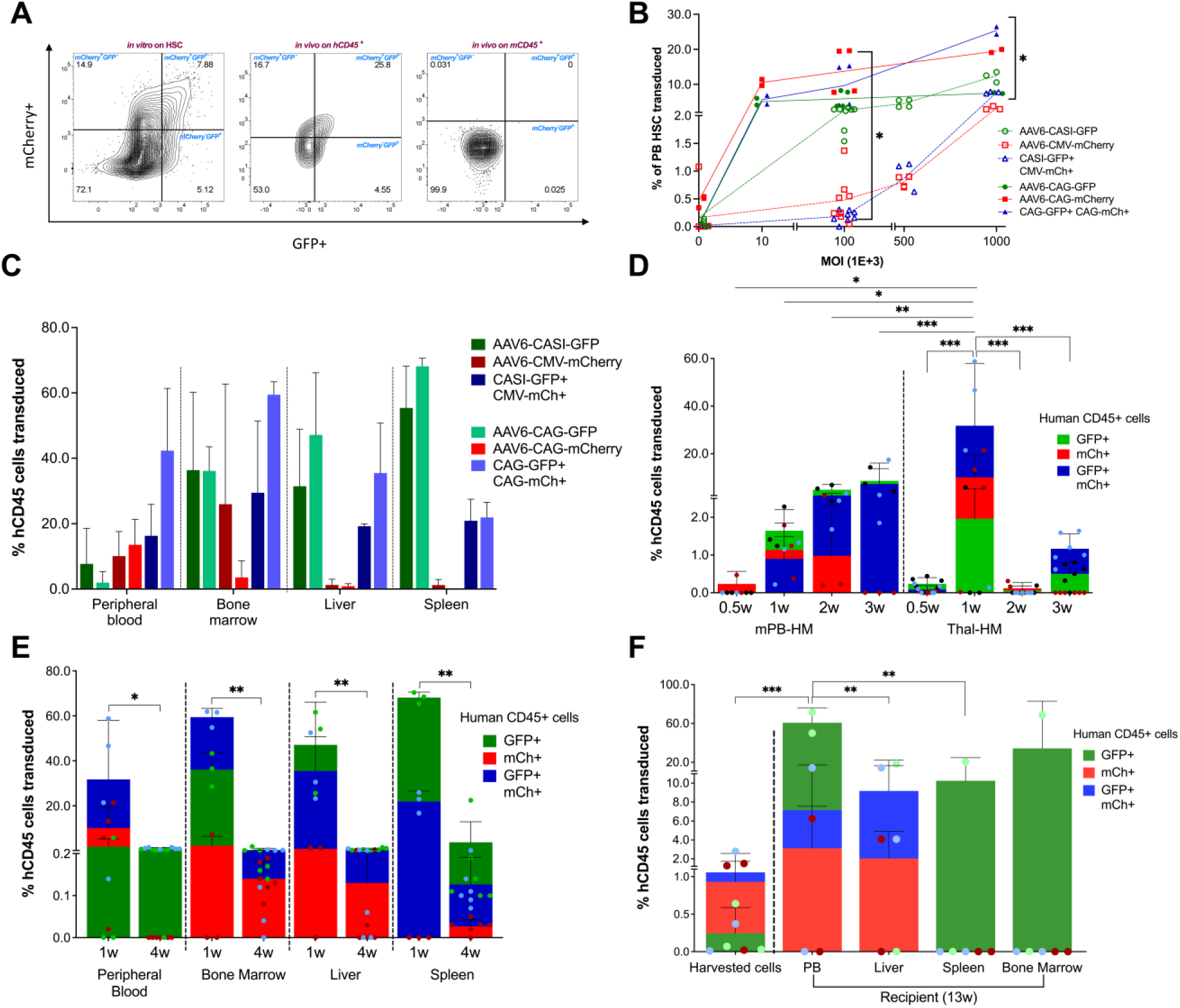
In vivo single and dual transduction of AAV6-GFP and AAV6-mCh in humanized mice. **A** Representative flow cytometry gating to assess single and dual transduction *in vitro* on HSC (left), and *in vivo* on human CD45^+^ (center) and mouse CD45.1^+^ (right). **B** In vitro PBHSC expression of either GFP or mCherry, or both, at varying MOI of each vector pair. **C** In vivo transduction of hCD45 cells recovered from mouse PB, bone marrow, liver and spleen expressing either GFP or mCherry, or both, following injection of both vector pairs at a dose of 5E+12vg/kg as a 1:1 mixture in Thal-HM (from one TDT patient donor) at 1w post-AAV injection. **D** In vivo single and dual transduction of hCD45 cells derived from PB of Thal-HM and mPB-HM given 5E+12vg/kg of AAV6-CASI-GFP and AAV6-CMV-mCherry mixture (1:1 ratio), at 0.5w– 3w post-AAV injection. **E** In vivo single and dual transduction of engrafted circulating and nested hCD45+ cells in Thal-HM (from three TDT patient donors) harvested 1w and 4w post-AAV injection. **F** Composition of transduced cells in retrieved “donor” hCD45 cells harvested from primary Thal-HM at 14w, and composition at harvest. *p<0.05, **p<0.01, ***p<0.001, ****p<0.0001.

**Figure 5.**
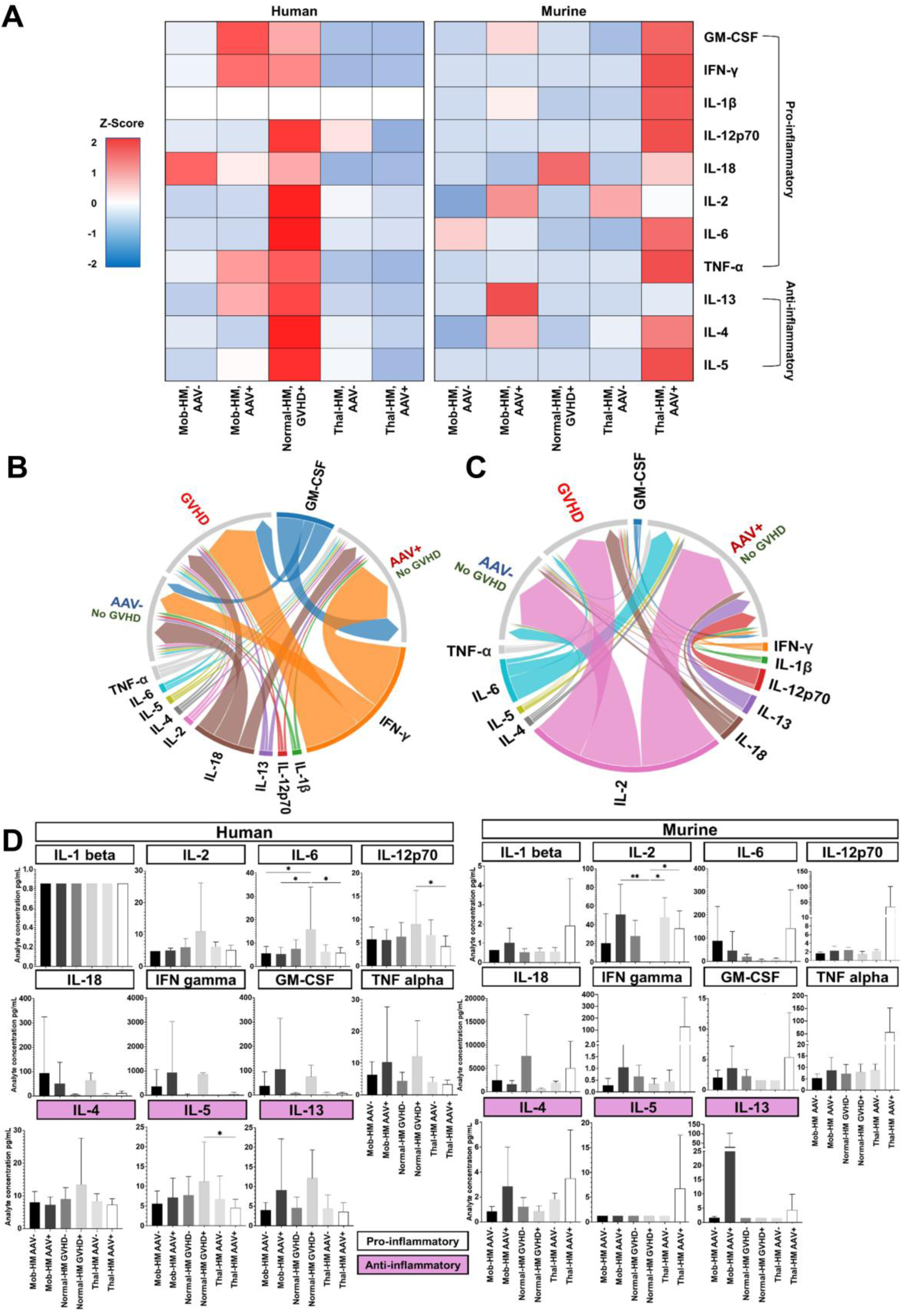
Humoral and cellular immune responses to transplanted cells and AAV6. **A** Heatmap showing human-and murine-origin cytokine levels in Normal-HM with graft-v-host disease (GVHD), mPB-HM and Thal-HM without GVHD, with and without AAV exposure, represented by Z-scores (±2SD from baseline cytokine levels measured in Normal-HM humice without GVHD). Chord diagrams representing fold-change of human (**B**) and murine (**C**) cytokine levels relative to Normal-HM without GVHD, in humice with GVHD, with and without AAV exposure (chord width represents fold-change). **D** Comparative expression of individual human and murine cytokines. *p<0.05, **p<0.01, ***p<0.001, ****p<0.0001. GVHD - graft-v-host disease.

We then compared single and dual transduction in circulating and nested hCD45 following *in vivo* AAV6 administration with both vector pairs. In Normal-HM, circulating hCD45 dual transduction was 16.30±9.59% (n=2) with CASI-GFP/CMV-mCh (single transduction ranged from 7.69-10.07%), while dual transduction was 42.31±19.07 (n=3) with CAG-GFP/CAG-mCh (single transduction 1.96-13.55%), with no significant differences (**Figure 4C**). There were also no differences in nested hCD45 transgene expression. Dual transduction with CASI-GFP/CMV-mCh was highest in BM at 29.49±21.94%, while single transduction ranged from 1.22±1.73% (mCh^+^) to 55.39±12.80% (GFP^+^), both in spleen (n=2). Dual transduction with CAG-GFP/CAG-mCh (n=3) ranged from 21.9±4.66% in spleen to 59.44±3.95% in BM, while single transduction ranged from 0.88±0.77% in liver (mCh^+^), to 68.14±2.48% in spleen (GFP^+^).

mPB-HM treated with CASI-GFP/CMV-mCh showed single transduction of circulating hCD45 reaching 1.14±0.71% mCh^+^ at 1w and 8.65±5.13% GFP^+^ at 3w, while dual transduction reached 7.35±8.77 at 3w (n=3, **Figure 4D**). We performed *in vivo* mobilization in a subgroup of mPB-HM with GCSF and AMD3100 (n=3), to interrogate this strategy of increasing availability of circulating hCD45 for transduction; PB hCD45 increased from 1.00±0.56% to 2.79±1.80% after 2w, returning to 0.29±0.82% in PB and 0.22±0.53% in BM by 4w post-mobilization (**Supplementary Figure S1**). GFP^+^ and GFP^+^mCh^+^ expression peaked at 18.35±13.56% and 22.31±26.87% respectively 3w post-injection, while mCh^+^ expression was 4.80±3.10% at 1w, and <1% at 3w. This upward trend following *in vivo* mobilization was not statistically significant, and the experiment was not repeated in other groups.

In serially-monitored Thal-HM (CASI-GFP/CMV-mCh, n=2; CAG-GFP/CAG-mCh, n=3, due to vector availability), single transduction in circulating hCD45 cells peaked 1w post-injection at 1.96±3.39% GFP^+^ (n=3) and 10.17±9.28% mCh^+^ (n=4), with GFP^+^mCh^+^ expression reaching 31.77±26.21% (n=4), significantly higher than single or dual transduction at other timepoints (p=0.01 to p<0.0005, **Figure 4D**). By 3w post-injection, transduction had declined to 0.50±0.30% GFP^+^ and 1.17±0.40% GFP^+^mCh^+^ (n=6).

In nested hCD45 (BM, liver and spleen), single transduction ranged from 0.88-2.38% (mCh^+^) to 36.13-68.14% (GFP^+^); GFP^+^mCh+ expression reached 21.91±4.66% and 59.44±3.95% in spleen and BM, respectively (**Figure 4E**). hCD45 in spleen maintained ∼4% GFP^+^ expression by 4w, while in other organs, single or dual transduction had dropped significantly to <1% (p<0.001, n=6). In contrast, murine mCD45 transduction was significantly lower than hCD45 transduction, with maximum single or dual transduction of ∼2% in liver murine mCD45 cells at 1w, and <1% in mCD45 cells from other organs (data not shown).

Transplantation of transduced hCD45 recovered from primary Thal-HM (0.25±0.34% GFP^+^, 0.93±0.80% mCh^+^ and 1.06±1.51% GFP^+^mCh^+^) resulted in persistent transduced cells in secondary recipients, which demonstrated expression levels up to 9.19-60.72% GFP^+^, 2.04-3.13% mCh^+^, and 7.15-9.19% GFP^+^mCh^+^ in circulating and nested hCD45. Circulating PB hCD45 showed higher GFP^+^ expression (60.72±15.15%) in secondary compared to primary recipients (0.25±0.34%, p<0.001). This was also significantly higher than in nested hCD45 GFP^+^ in liver (9.19±12.99%) and spleen (10.26±14.50%, **Figure 4F**).

### Human and murine humoral and cellular responses to *in vivo* AAV6

Murine-and human-origin cytokine expression in Thal-HM (given AAV^+^ or not given AAV^-^), mPB-HM (AAV^+^ or AAV^-^) and Normal-HM with GVHD were compared against baseline expression in Normal-HM without GVHD or AAV exposure. Z-scores (±2SD from baseline levels in Normal-HM humice without GVHD) shown in the heatmap indicate that the highest expression of all human pro-inflammatory and anti-inflammatory cytokines were observed in Normal-HM with GVHD. Thal-HM (AAV^+^ or AAV^-^) had Z-scores of 0 to-2 for all human cytokines, while mPB-HM-AAV^+^ showed increased human GM-CSF, IFN-γ, TNF-α and IL-13 expression following AAV exposure (**Figure 5A**). The highest Z-scores of murine cytokines were observed in Thal-HM-AAV^+^, with more limited positive responses in mPB-HM-AAV^+^; murine cytokines were minimally affected in GVHD mice.

Of all human cytokines, the greatest upregulation was observed in pro-inflammatory IFN-γ, GM-CSF and IL-18 (shown by the widest chords in the chord plot) expressed by all AAV^+^ and GVHD humice (**Figure 5B**). Among murine cytokines, IL-2 was upregulated in all three groups, while AAV exposure elicited more robust murine IL-6, IL-13, IL-12p70 expression, and GVHD humice produced marginally higher IL-18 (**Figure 5C**). Larger analyte differences were observed among murine anti-and pro-inflammatory cytokines than among human cytokines. GVHD humice expressed significantly higher human IL-6, IL-12p70 and IL-5 than observed in mPB-HM (AAV^+^ and AAV^-^) and Thal-HM (AAV^+^), while IL-1β was below detection threshold in all groups (**Figure 5D**). Human anti-AAV6 antibodies were not detected in humice at sacrifice (**Supplementary Figure S4**).

## DISCUSSION

Our model demonstrates the feasibility of *in vivo* dual AAV6-mediated transduction of human HSC-derived hCD45 with and without β-thalassemia mutations, showcasing a strategy that can be used to deliver large GMT tools using a split-intein technique. Though the techniques utilized are not novel, as we used marker transgenes, our objective was to build this patient-specific model to eventually interrogate novel gene and base editors *in vivo*. Humanized mouse models, mostly using HSC from healthy donors, UCB or occasionally patients with sickle cell disease, have been described, but published models specifically using HSC from transfusion-dependent β-thalassaemia patients are rare. To our knowledge, a patient-derived humanized model specifically for *in vivo* gene therapy with non-integrating AAV6 vectors has not been previously described. Our model offers a novel strategy towards adopting a personalised therapeutic approach to individuals with β-haemoglobinopathies, that is relatively simple to produce (peripheral mobilisation is not required), permits rigorous assessment of multiple outcomes in vivo (including toxicity and long-term survival of transduced patient-derived cells), and uncovers individual barriers to in vivo GMT (e.g. low targeting efficiency).

Our model achieves similar chimerism and transgene expression as humanized models reconstituted with healthy donor HSC, using helper-dependent adenovirus and in vivo selection to enrich for transduced HSC;^30^ in contrast to “normal” models, our approach recapitulates the biology and genetic complexity of the intended recipients of these therapies, while direct in vivo evaluation of diseased HSC provides critical feasibility and safety data that can potentially accelerate translation to early-phase in vivo clinical trials. With such a system, we can produce a range of patient-specific editing tools targeting the common β-globin mutations for *in vivo* use, and test efficacy and precision with a relatively low-cost rapidly-producible model (**Figure 6**). Further benefits include enhanced clinical decision-making by identifying patients most likely to benefit from in vivo gene therapy while flagging potential complications before treatment initiation.

**Figure 6.**
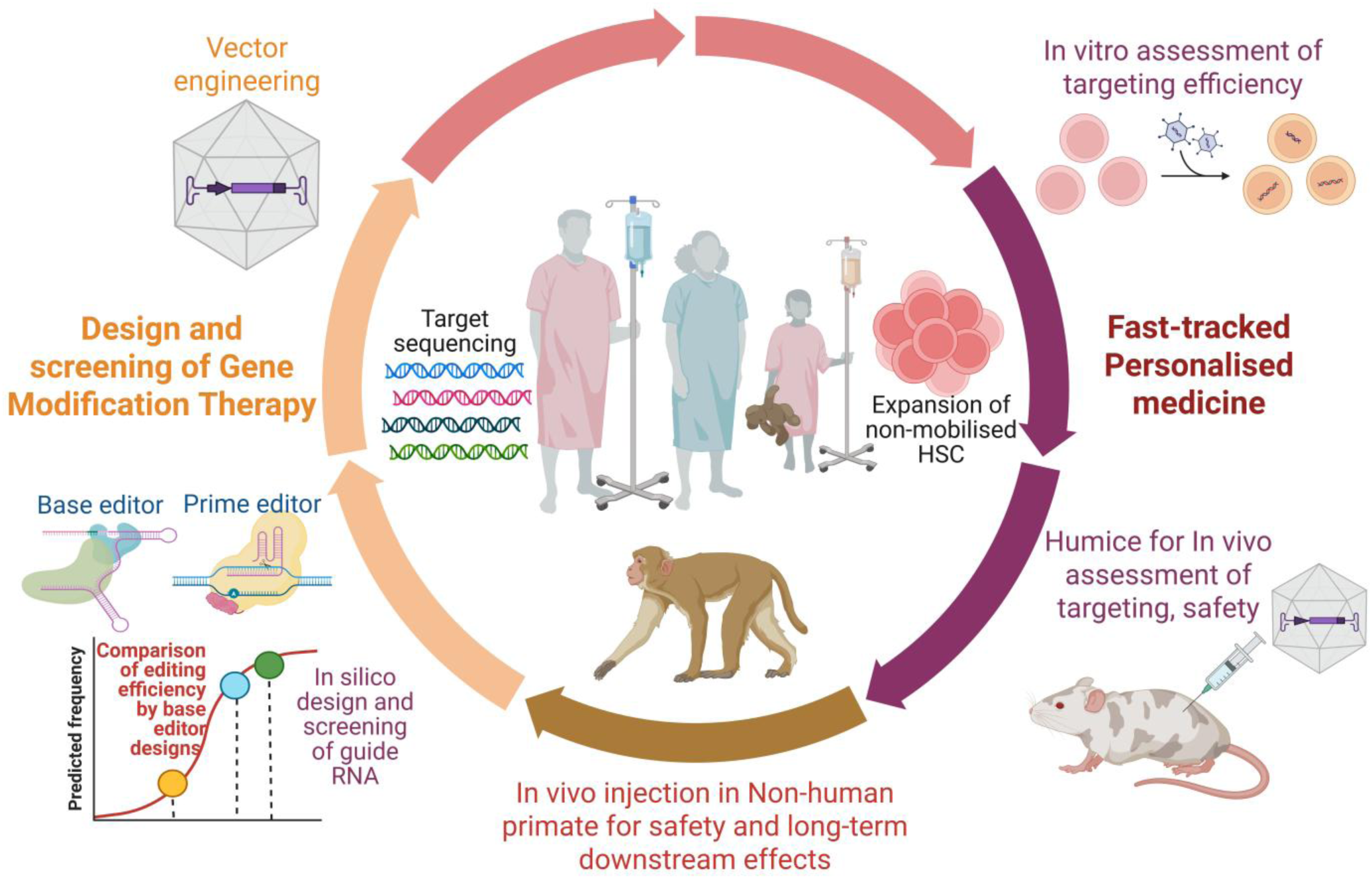
A proposed clinical translational pathway towards *in vivo* gene modification therapy for thalassaemia. Individual patient mutations are sequenced and editing tools, comprising guide RNA and appropriate base or prime editor, are designed and screened in silico for predicted targeting efficiency. Effective gRNA-editor are packaged into an appropriate vector, screened *in vitro* and *in vivo* (in patient-derived humice) for editing frequency and other cellular effects. The appropriate vector can additionally be applied directly in a non-human primate to elicit clinically useful data on toxicity and durability of effect. This strategy can be applied to individual mutations causing transfusion-dependent thalassaemia, and can be extended to primary immunodeficiencies, non-deletional α-thalassaemia and other genetic causes of bone marrow failure. Created in https://BioRender.com

1. Humanized mice can be reconstituted with non-mobilized expanded peripheral blood HSC from adults and children with TDT

Humice are typically reconstituted with human umbilical cord blood (UCB) HSC (comprising 0.3-2% CD34^+^) or with mobilised PBHSC (0.1-2% CD34^+^ cells) and can achieve human chimerism of 1-5%.^31, 32^ We and others have described humice reconstituted with non-mobilised PBHSC,^33^ and we have utilised *in vitro* expansion to enhance chimerism, which is directly related to donor inoculum quantity. Our model serves as a relatively simple method for producing patient-derived humice from a limited quantity of adult HSC derived from regular venepuncture samples without mobilisation, avoiding potential risks of splenic rupture in TDT, and providing a more accessible platform for high-throughput *in vivo* screening of novel HSC-targeting gene therapies.^34^ As 10-40% of donors are “poor mobilizers”, unable to release a sufficient number of nested HSC into circulation, this strategy is useful for personalised testing of new GMTs in an informative model.^19^ ^34^

Expansion of non-mobilised PBHSC from TDT and healthy volunteers resulted in 3-4-fold increase in cell counts while retaining >50% CD34^+^ cells, with ST-HSC and LT-HSC ranging from 2.5-12.5%, and achieved reasonable donor chimerism of 4-10% by 12-14w post-transplantation in healthy NSG mice with no signs of GVHD, despite the restricted CD34^+^ quantity of donor cells. Secondary transplantation confirmed the presence of long-term repopulating hCD45, though our model demonstrated mostly myeloid reconstitution; additionally, the increased proportion (∼60%) of single-and dual-transduced engrafted hCD45 suggest at least a survival benefit and absence of significant cell toxicity, and lends support to the proliferative advantage acquired by modified cells, as reported by others.^35^ The ability to maintain long-term HSC repopulation post-transduction highlights the possibility of a one-time curative treatment for the β-haemoglobinopathies, and supports the use of a dual-AAV6 approach to edit HSC *in vivo* without need of *ex vivo* manipulation or myeloablation.

The absence of hCD34^+^ cells may be related to transgenic overexpression of human cytokines causing HSC exhaustion, a reported limitation of long-term CD34^+^ engraftment in humanized models.^36^ Robust expansion of TDT-patient primary HSC may reflect greater propensity towards cell division and replication due to increased susceptibility to haemolysis.^24^ Robust expansion of TDT-patient primary HSC may reflect greater propensity towards cell division and replication due to increased susceptibility to haemolysis.^24^ In comparison, our humice reconstituted with commercially available GCSF-mobilised PBHSC, comprising 76% CD34^+^ cells, attained similar chimerism of 3-9% for a longer duration of 20w post-transplantation. Foetal and UCB-HSC achieve greater engraftment efficacy and transducibility due to the higher concentration of CD34^+^CD133^+^ HSC.^37, 38^ As both mobilised and non-mobilised adult PBHSC achieved similar chimerism levels in our humice, we infer that donor cell immaturity or “stemness” is the more critical factor producing high chimerism in humice. We did not transplant UCB-HSC as putative recipients of *in vivo* GMT would be infants or adults rather than neonates, thus *in vivo* transduction of adult HSC was the research aim.

2. Personalized humice adds value to clinical trials by permitting prediction of targeting efficiency and immune responsiveness to candidate vectors

Personalised humice provide direct insight of gene modification on the differentiation, behaviour and interaction of human HSC within hematopoietic niches, allowing personalised assessments of efficacy and safety, immune responses and direct cell toxicity, and permit analyses of delivery or enhancement techniques that may be ethically-challenging to perform in patients, aiding in the clinical approval of novel GMTs.^35^ Clinically relevant readouts include frequency of correct *in vivo* editing, insertions/deletions and incorrect or off-target editing, persistence of transgene expression, vector integration, and humoral and cellular responses to vector, transgene or editor, and the ability to target long-term repopulating cells. This data is informative in predicting the individual’s response to *in vivo* GMT. Furthermore, we demonstrated both single and dual *in vivo* transduction of human HSC with AAV6, from both TDT and non-thalassemic patients, attesting to the value of this model for developing AAV-GMTs targeting HSC; this model is particularly useful as *in vivo* AAV6 transduction is not well-demonstrated in murine haemoglobinopathy models due to the low AAV transduction efficiency of normal and abnormal murine HSC.^23, 39^ We showed distinct profiles of human and murine CD45^+^ cell transduction (representing erythroid and lymphoid lineages) and of human and murine cellular responses in this model, allowing us to reliably assess human target cell behavior *in vivo*. Limitations of this model include low hCD34^+^ chimerism restricting the ability to perform downstream analyses on human hematopoiesis, development of GVHD, deficiencies in other human blood components, and low-fidelity human cell behavior and development due to the lack of species-specific factors.^35^

3. Potential advantages of *in vivo* AAV6-GMT for haemoglobinopathies and other HSC diseases

*Ex vivo* gene modification and autologous transplantation require myeloablation and conditioning which increases therapy-related complication risks of opportunistic infections and malignant transformation.^9^ Recombinant and engineered AAVs are currently leading the *in vivo* delivery platform development and are in clinical trials for wide-ranging applications due to their low pathogenicity, broad tissue tropism and relatively high delivery efficacy.^40^ With gene addition technologies, transient expression is expected, rendering this unsuitable for haemoglobinopathies in which lifelong production of normal globins is necessary; with editing strategies however, transient episomal appearance of the non-integrated AAV is not a hindrance if *in situ* gene correction can occur efficiently, particularly if survival and proliferation advantage of corrected HSC is conferred.^41^

Additional benefits of gene editing over transgene addition, particularly in the context of transgene integration required for haemoglobinopathies and primary immunodeficiencies, include not disrupting neighboring regulatory genes, and lower risks of off-target effects in non-hematopoietic cells.^42^ Achieving this with *in vivo* administration may further reduce the physical, procedural, healthcare resource and socioeconomic burdens of therapy, by simplification of processes and minimization of concomitant conditioning.^21, 43^

The innate resistance of HSC to gene modification, particularly by non-integrating and non-viral vectors, is a challenge;^8, 18^ additionally, true long-term repopulating stem and progenitor cells are rare in adult HSC (<0.01% in adult peripheral blood) and show variable transducibility due to population heterogeneity.^37^ Gene-modified HSC lose their repopulating and engraftment capacity, particularly after *ex vivo* culture and cryopreservation.^44^ These factors contribute to subtherapeutic gene modification of circulating PBHSC, which could be overcome by mobilising bone marrow HSC prior to therapeutic injection, or by utilising intraosseous delivery.^18, 45^ ^37^

HSC-directed GMT currently follows complex protocols of *ex vivo* genetic modification by viral-mediated or physical means of cell entry (the latter of which may cause cellular damage^46^), cell product screening for potentially oncogenic off-target mutations (to minimize clonal expansion of mutant HSC) followed by autologous transplantation, together with myeloablative pre-conditioning (which may cause pancytopenia and opportunistic infections) to prepare the hyperplastic bone marrow for engraftment.^12^ The move towards *in vivo* GMT for HSC diseases will substantially reduce these physiological burdens and medical expenditures related to production and patient interventions.^47^

4. *In vivo* single or dual AAV transduction strategies targeting HSC have wide applicability for gene therapies

*In vivo* AAV-mediated GMT is the most widely employed therapeutic strategy, available commercially for a range of genetic conditions, from rare lysosomal storage diseases and congenital blindness,^3, 4, 48^ to the hemophilia, spinal muscular atrophy, Duchenne muscular dystrophy and cystic fibrosis.^49–52^ Therapeutic products are delivered directly into the target tissues (intraocular or intrathecal) or systemically (intravascular or intramuscular), relying on the widespread biodistribution of AAV and specific tropism of various serotypes for the cells of interest.^14^ The timing of GMT administration is critical to optimizing outcomes, particularly in thalassemia, as intervention early in the development spectrum ensures more efficient HSC targeting, when the quantity of circulating CD34+ HSC is highest and when HSC are concentrated in the liver.^38^ Direct intrahepatic injections or systemic delivery methods can be strategically employed to ensure a higher density of HSCs is targeted. In vivo editing is accelerating towards clinical applications, and fast-tracked N=1 in vivo trials may become more common, as demonstrated with hepatocyte-directed in vivo base editing of the pathogenic *CPS1* variant in a young infant using a customized adenine base editor, highlighting the potential of this approach for personalized gene therapy.^53^

AAV6 demonstrates tropism for HSC *in vitro* and *in vivo*.^54, 55^ While single-and dual-AAV6 transduction occurred in 5.7±6.2% of PBHSC *in vitro*, engrafted thalassemia and non-thalassemia hCD45 demonstrated single-AAV6 transduction in 1.7±3.8% of circulating hCD45 (mean PB chimerism of 13.5±20.2% at the time of assessment), and a robust dual-AAV6 transduction of 21.9±3.8%–59.4±3.2% in nested hCD45 by 1w post-administration, particularly in liver and spleen. The higher concentration of transduced cells nested in solid organs may be the result of active migration and homing into haemopoietic and reticulo-endothelial niches, or possibly of active uptake of AAV6 by hCD45 in these niches.^56^ This may reflect synergistic microenvironmental and niche factors and active processes that enhance AAV6 cell entry and intracellular trafficking.^8, 39, 56^ We have emphasized dual-AAV6 transduction as split-intein editor design has been used effectively to correct murine models of genetic deafness, amyotrophic lateral sclerosis, neurodegenerative ataxia and inborn errors of hepatocyte metabolism, effectively transducing murine brain, liver, retina, heart and skeletal muscle.^57, 58^ A clinical trial is underway to using AAV-dual technology to treat retinitis pigmentosa related to Usher Syndrome Type 1B.^59^ This strategy may shorten the runway to clinical translation across the various HSC genetic diseases resulting in impaired hematopoiesis (β-haemoglobinopathies) or immunity (severe primary immunodeficiency syndromes), fetal mortality (α-thalassemia major), malignancies or bone marrow failure,^1, 2, 60^ not all of which are curable by allogenic HSCT.^60 57, 58^

High VCN (19257.65-84541.42 copies/cell) in engrafted Thal-PBHSC compared to the substantially lower VCN (0.04-3810.93 copies/cell) in non-thalassemia PBHSC may possibly suggest greater susceptibility of thalassemia HSC to vector entry. VCN in IVS1-5G>C cells from an adult TDT patient was substantially higher than in IVS2-654C>T cells donated by child patients, corresponding to higher transgene expression in the former; while we cannot draw conclusions on age and mutation affecting HSC transducibility, we acknowledge that these may be two of several biological or interventional factors affecting *in vivo* transduction, regardless of vector dose.^61^ Despite this, the milder immune responses we observed are encouraging and in line with reported safety of AAV. The potentially large variability between patients with the same disease reinforces the value of a personalized model for *in vivo* testing of novel GMTs, as we present here.

## CONCLUSION

We have demonstrated the feasibility of *in vivo* transduction of human adult HSC with a single-or dual-AAV6 strategy in patient-specific humice. We report generally high levels of transduction in circulating and nested HSC which did not produce clinically significant immunotoxicity, though varying degrees of cellular responses were observed. Engrafted thalassemia HSC displayed more robust human cytokine expression than non-thalassemia HSC in response to AAV6. This is a clinically relevant *in vivo* model for HSC-targeting therapies that can be utilized with relative ease to assess personalized cellular effects and potential immune responses. *In vivo* genetic editing strategies may be ideal for HSC diseases if off-target effects are minimized, particularly with emerging base and prime editors, and novel nucleases designed to enhance precise correction. Dual-AAV6 transduction can accommodate large editors via split-intein design, and the high but transient transduction we observed is useful, allowing sufficient time to facilitate editing with rapid cell clearance to minimize cell toxicity.

## Abbreviations

PB: peripheral blood
BM: Bone marrow
HSC: hematopoietic stem cells
PBHSC: peripheral blood HSC
mPBHSC: mobilized PBHSC
HSPC: hematopoietic stem and progenitor cells
UCB: umbilical cord blood
LT-HSC: Long-term HSC
ST-HSC: short-term HSC
AAV: adeno-associated viral
CRISPR: clustered regularly interspaced short palindromic repeats
GMTs: Gene modification therapies
TDT: transfusion-dependent β-thalassemia
GVHD: graft-v-host disease
cGMP: Current good manufacturing practices
GCSF: granulocyte colony stimulating growth factor
BFU-E: Burst-forming unit-erythroid
CFU-E: Colony-forming unit-erythroid
CFU-GEMM: colony-forming unit-granulocytes/erythroids/macrophages/megakaryocytes
NSG: NOD.Cg-Prkdcscid Il2rgtm1Wjl/SzJ
Normal-HM: humanized mice engrafted with expanded non-thalassemia HSC
Thal-HM: humanized mice engrafted with expanded thalassemia HSC
mPB-HM: humanized mice engrafted with mobilized, non-expanded non-thalassemia HSC
VCN: vector copy number
IFN-γ: Interferon gamma
TNF-α: Tumor necrosis factor-alpha
IL: Interleukin

## Conflict of Interests

The authors declared that they have no conflict of interests.

## Ethics Approval and Consent to Participate

Ethics approvals were obtained from the National University of Singapore Institutional Review Board (NUS-IRB-2021-424), and the National Healthcare Group Domain-Specific Research Board Domain D (NHG DSRB 2021/00783). Mice were bred and housed under specific pathogen-free conditions at the Biological Resource Center (BRC) of the Agency for Science, Technology and Research (A*STAR), Singapore. All animal experiments conformed to the Animal Research: Reporting of *In vivo* Experiments (ARRIVE) guidelines, and were conducted in strict accordance with the guidelines for the Care and Use of Animals for Scientific Purposes, which are released by the National Advisory Committee on Laboratory Animal Research, Agri-Food & Veterinary Authority of Singapore, and the International Animal Care and Use Committee (IACUC) at A*STAR which specifically approved this study (IACUC 221686).

## Notes

**Acknowledgements:** This study was funded by the Singapore Ministry of Health National Medical Research Council NMRC/CSA INV/0012/2016 and NMRC/CSAINV20nov-0016. CNZM is supported by grants from the NMRC (NMRC/TA/0003/2012, NMRC/CSA-INV/0012/2016, and NMRC/CSAINV20nov-0016). The funding body played no role in the design of the study and collection, analysis, and interpretation of data and in writing the manuscript. Multiplex measurements were carried out at the Stem Cell Core Facility supported by the Healthy Longevity Translational Research Programme at the Yong Loo Lin School of Medicine, National University of Singapore. We thank Daryl Lim (Genome Institute of Singapore), Nuryanti Johana, Lay-Geok Tan and Sabrina Adam (National University of Singapore) for their technical assistance. Artificial intelligence (AI) was employed to assist in the formulation and refinement of the language and conceptual development presented.

### Competing Interest Statement

The authors have declared no competing interest.

