## Supplemental for "Modeling Adeno-Associated Viral Vector 6-mediated *In Vivo* Gene Delivery to Expanded Non-Mobilized Haemopoietic Stem Cells from Transfusion-dependent Thalassemia Patients in a Humanized Mouse"

This file includes Supplemental Results, Supplemental Figures and Figure Legends, and Supplemental Tables.

**Supplemental Results**

We evaluated vector concentration in circulating and nested hCD45 at terminal harvest. A wide range of vector copy number per cell (VCN) in Thal-HM and mPB-HM was detected, from 19257.65±39398.00 vs. 176.72±185.83 in BM, 26844.78±34626.06 vs. 3810.93±4024.08 in liver and 84541.42±130.49 vs 0.04 in spleen, respectively (**Supplemental Figure S3A**). We found higher VCN in nested hCD45 carrying IVS1-5G>C mutation than in cells bearing IVS2-654C>T in mouse spleen, correlating with the higher transgene expression in Thal1-HM (202651.80±130127.37 vs. 176.86±217.84, $p<0.0001$, (**Supplemental Figure S3B**).

**Supplemental Figures**

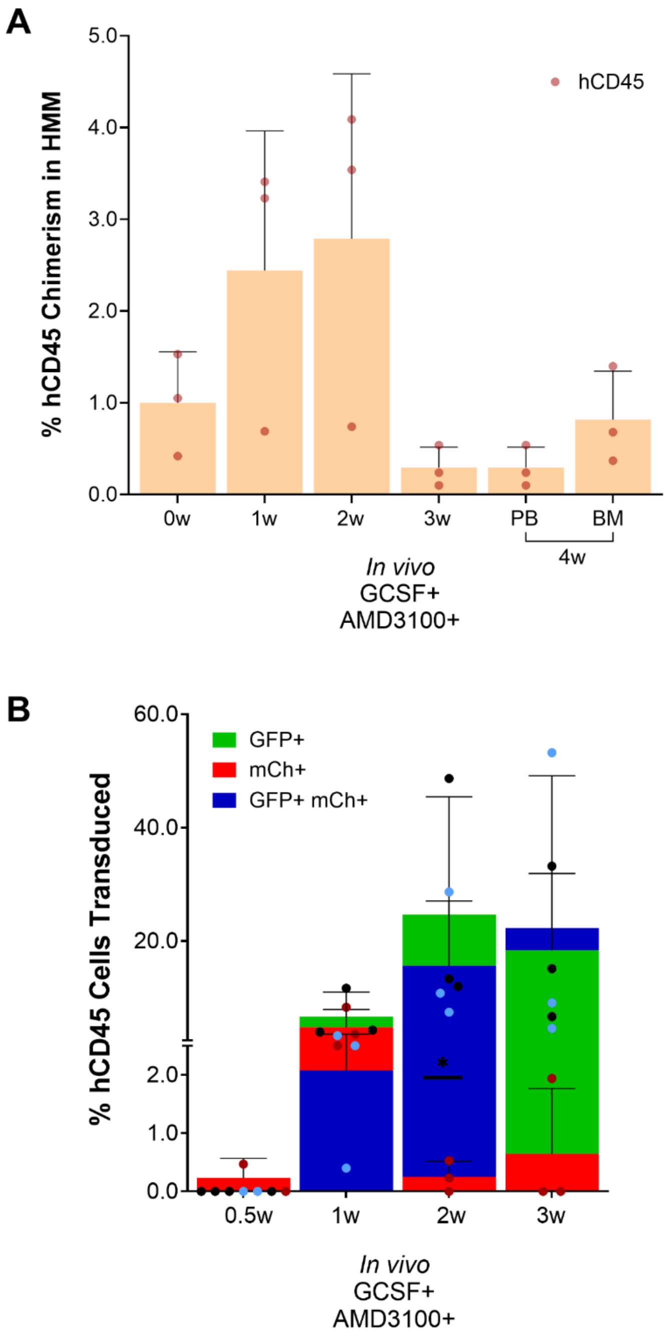


**Figure S1.** ***In vivo mobilization of hCD45 cells in Humanized Mice.*** Granulocyte-Colony Stimulating Factor (GCSF) and AMD3100 injections were given to Mob-HM mice to mobilize hCD45 cells homed in the bone marrow. **A** Percentage of hCD45 cells in PB 0 – 3 weeks post injection, and in PB and BM at time of harvest (4 weeks post injection). **B** *In vivo* single and dual transduction of hCD45 cells derived from PB of GCSF and AMD3100 injected Mob-HM mice, that were given 5E+12vg/kg AAV6-CASI-GFP and AAV6-CMV-mCherry mixture (1:1), at 0.5w–3w post-AAV injection.

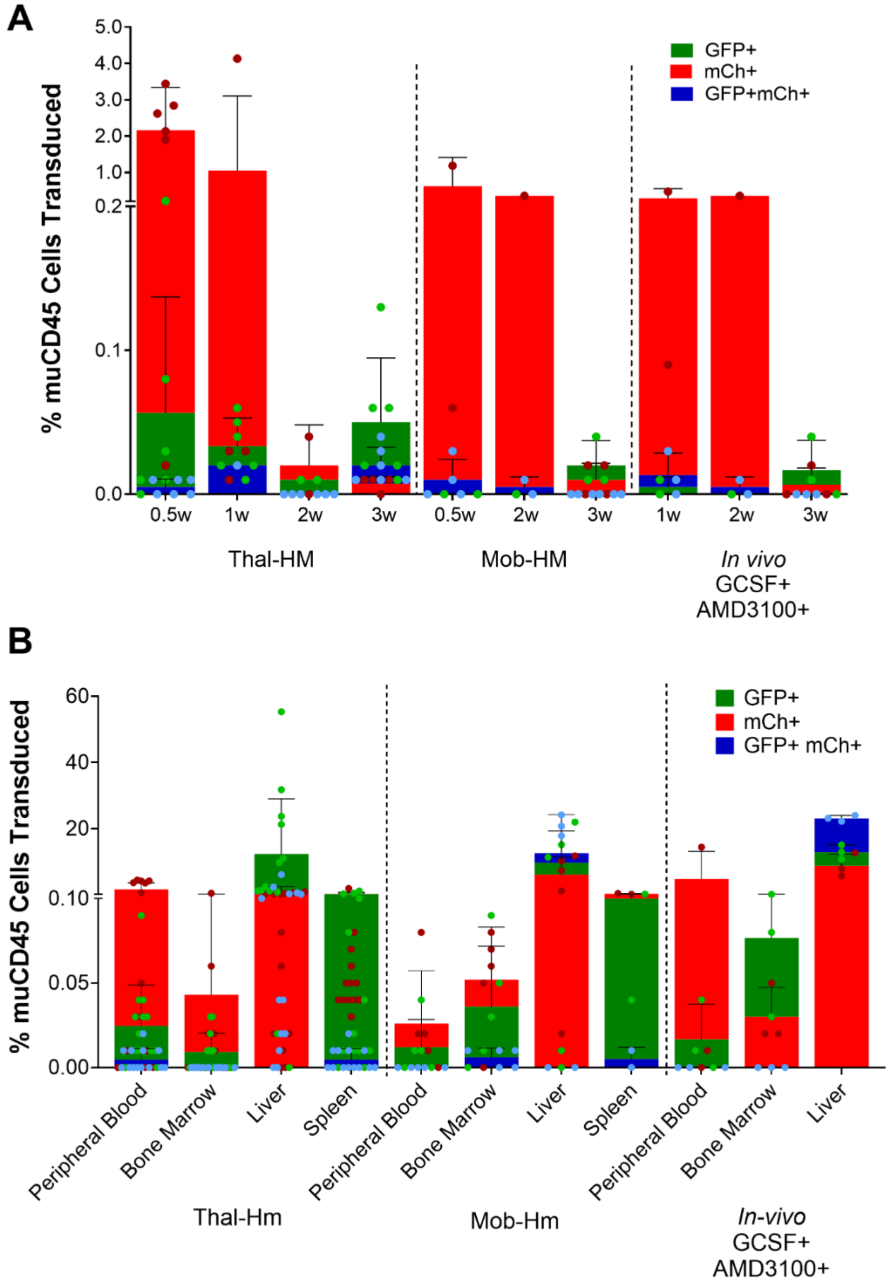


**Figure S2.** ***In vivo single and dual transduction of AAV6-GFP and AAV6-mCh in Murine CD45⁺ (mCD45⁺) cells of Humanized mice.*** **A** *In vivo* single and dual transduction of mCD45+ cells derived from PB of Thal-HM, Mob-HM, and GCSF and AMD3100 injected Mob-HM that were given 5E+12vg/kg AAV6-CASI-GFP and AAV6-CMV-mCherry mixture (1:1), at 0.5w–3w post-AAV injection. **B** *In vivo* single and dual transduction of mCD45+ cells derived from harvested tissues of Thal-HM, Mob-HM, and GCSF and AMD3100 injected Mob-HM that were given 5E+12vg/kg AAV6-CASI-GFP and AAV6-CMV-mCherry mixture (1:1).

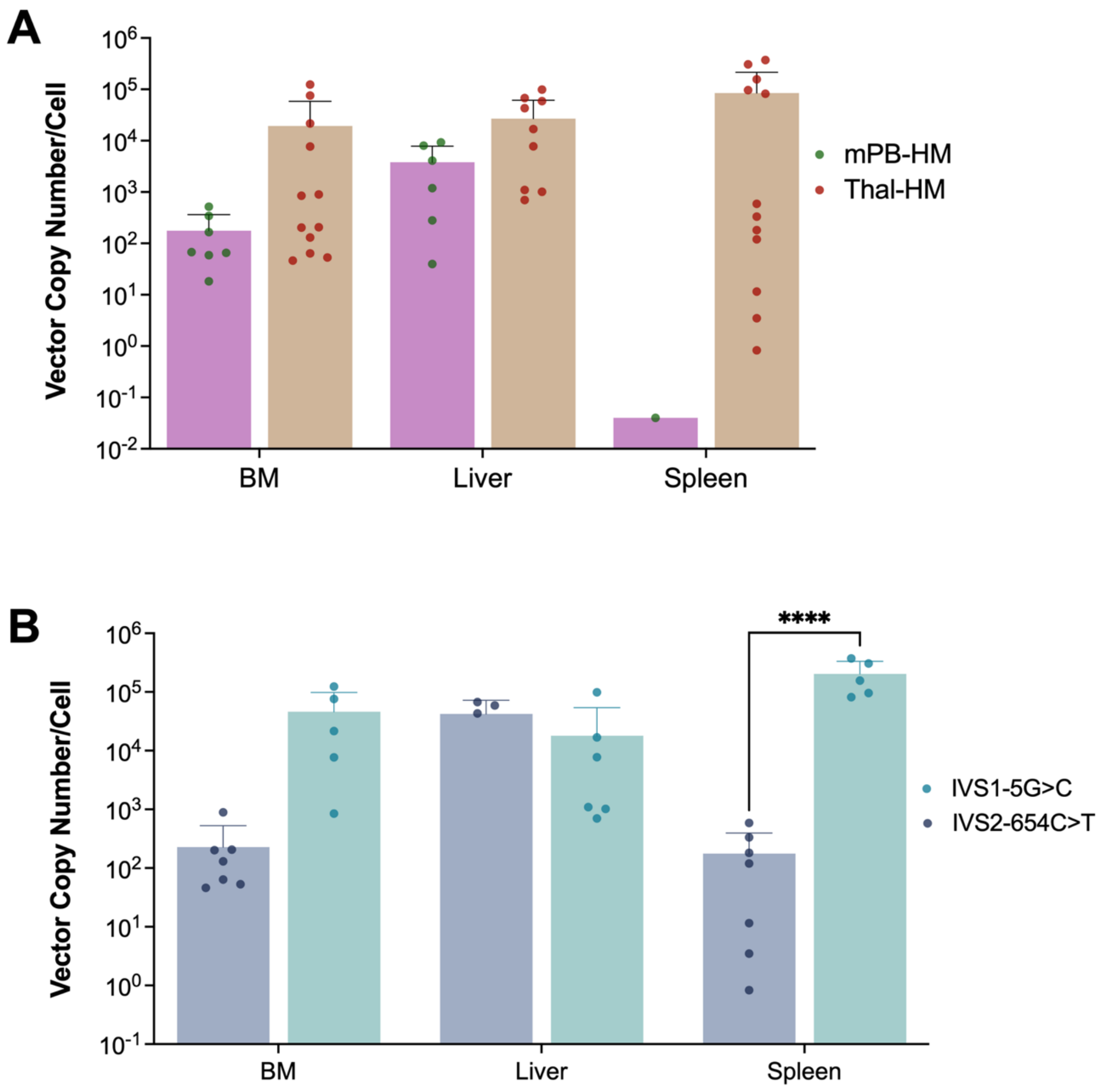


**Figure S3. Efficacy of Transgene Expression. A** Vector copy number in circulating and nested hCD45 cells in BM, liver and spleen cells in mPB-HM and Thal-HM. **B** Vector copy number analysis within Thal-HM group with IVS1-5G>C and IVS2-654C>T mutation. ****$p < 0.0001$

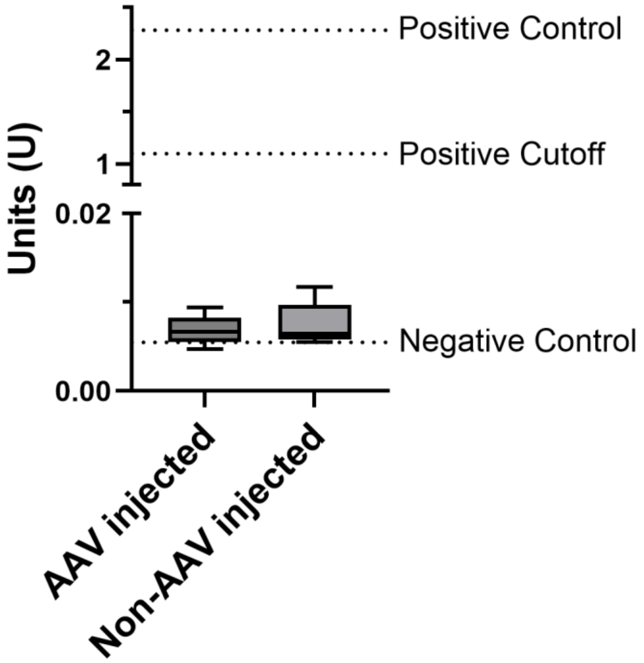


**Figure S4. Human Anti-AAV6 Antibody Expression in Humice.** Absorbance values were measured at 450/620 nm. Mean absorbance (OD) was calculated, with results expressed in arbitrary units (U) using the formula: Units (U) = (Avg OD) / (Cut-off), where cut-off value is based on Avg OD of manufacturer-provided cut-off control, measured here at 1.276. Human anti-AAV6 was not detected in all samples, with values close to negative control reading in both AAV injected (n=15) and non-AAV injected (n=6) groups.

**Supplementary Table**

Table S1. *Flowcytometry Antibody*

| HSC Characterization | | | |
|---|---|---|---|
| **Ab** | **Clone** | **Fluorescence** | **Brand** |
| Sytox Blue (Live/Dead) | - | BV-421 | eBioscience, Invitrogen |
| Lineage (CD2, CD3, CD14, CD16, CD19, CD56, CD235a) | RPA-2.10, OKT3, 61D3, CB16, HIB19, TULY56, HIR2 | FITC | eBioscience, Invitrogen |
| CD34 | 4H11 | APC | eBioscience, Invitrogen |
| CD133 | EMK08 | PE | eBioscience, Invitrogen |
| CD38 | HIT2 | PE-CF594 | Biolegend |
| CD90 | eBio5E10 | PE-Cy7 | eBioscience, Invitrogen |
| CD49f | GoH3 | BV-650 | BD Biosciences |

| HMM | | | |
|---|---|---|---|
| **Ab** | **Clone** | **Fluorescence** | **Brand** |
| Sytox Blue (Live/Dead) | - | BV-421 | eBioscience, Invitrogen |
| CD45 (Human) | HI30 | BV-711 | eBioscience, Invitrogen |
| CD45.1 (Mouse) | A20 | PE | eBioscience, Invitrogen |
| CD34 | 4H11 | APC | eBioscience, Invitrogen |
| CD3 | UCHT1 | PE-Cy7 | Biolegend |

Table S2. *PCR Primers*

| Target Sequence | Primer | |
|---|---|---|
| CAG Promoter | Forward | 5′-GTCAATGGGTGGAGTATTTACGG-3′ |
| | Reverse | 5′-AGGTCATGTACTGGGCATAATGC-3′ |
| AAV-ITR | Forward | 5′- GGAACCCCTAGTGATGGAGTT-3′ |
| | Reverse | 5′- CGGCCTCAGTGAGCGA -3′ |
| Human GAPDH | Forward | 5′- ACCCGTTGACTCCGACTT-3′ |
| | Reverse | 5′- ACCACTAGGCGCTCACTGTTC-3′ |

Table S3. *Procartaplex Target Proteins*

| Target Protein | Bead Region |
|---|---|
| GM-CSF | 44 |
| IFN-γ | 43 |
| IL-1β | 18 |
| IL-2 | 19 |
| IL-4 | 20 |
| IL-5 | 21 |
| IL-6 | 25 |
| IL-12p70 | 34 |
| IL-13 | 35 |
| IL-18 | 66 |
| TNF-α | 45 |
